# Experience and relatedness influence mating interactions in a simultaneously hermaphroditic snail, *Physa gyrina*

**DOI:** 10.1101/865170

**Authors:** Thomas M. McCarthy

**Affiliations:** Department of Biology, Utica College, Utica NY, 13502, U.S.A.

## Abstract

The means by which animals assess potential mates is an important issue in studies of reproductive systems. I tested whether an individual’s previous experiences and the relatedness of mates affected mating behavior in a simultaneous hermaphrodite snail, *Physa gyrina*. Previous work with this species showed reduced reproductive success resulting from both strong outbreeding and inbreeding. Thus, I predicted that individuals should prefer partners of intermediate relatedness. During activity trials, snails moved longer distances when exposed to chemical cues from conspecifics of lesser relatedness. Furthermore, during mating interactions, behavioral responses to relatedness varied with gender-role: male-role behaviors did not vary across relatedness treatments, while snails paired with either closely related or highly dissimilar partners increased their female-role resistance behaviors as interactions escalated. Experiences with their current partner also affected behavioral dynamics. Familiar pairs had fewer matings and longer latency times until a mating occurred than unfamiliar pairs. Snails acting in the female role also exhibited higher resistance rates in familiar pairs than in unfamiliar pairs. Previous, brief exposure to chemical cues in a non-mating context also influenced behavior during a subsequent mating interaction. Snails that were previously exposed to chemical cues from unfamiliar individuals tended to be more likely to occupy the male role following an encounter, and had significantly lower copulation frequencies and higher female-role resistance rates (i.e. were choosier) than those previously exposed to cues from familiar individuals. Overall, the results show that: 1) relatedness, past exposure to conspecific chemical cues, and experience with a current partner all influence mating behaviors in these snails; and 2) in these simultaneous hermaphrodites, an individual’s responses depend on whether it is occupying the male or female role.

## Introduction

A major focus in behavioral ecology is the study of mate choice. Considerable effort has been dedicated to studying kin recognition, its mechanisms and its implications [see 1–3]. Many studies have also examined the fitness effects of inbreeding and outbreeding [4–6]. Most animals prefer some potential mates to others and do not mate indiscriminately [7–9]. Consequently, mating behaviors should reflect fitness patterns [10–12]. One way to minimize the deleterious fitness effects of inbreeding and outbreeding is to be able to recognize kin [10] and alter mating behaviors accordingly. However, individuals’ mating behaviors and mate-choice decisions may often be influenced, sometimes in disparate ways, by their previous experiences (e.g. exposure to phenotypes during development, or previous social and mating interactions as adults) [13].

Genetic similarity of mates may dramatically affect fitness through changes in the number, viability or quality of offspring [11,14,15]. Optimal outbreeding can be viewed as a trade-off between conflicting evolutionary and selective forces [5], i.e. inbreeding and outbreeding. Inbreeding occurs when related individuals produce offspring [16], or when hermaphrodites self-fertilize. Individuals may often encounter relatives as potential mates [10,11,17]. Conversely, outbreeding occurs between unrelated individuals within a population, or between individuals originating from different populations following a migration event. Both inbreeding and outbreeding have varying effects on reproductive success. If inbreeding and outbreeding both result in depressed fitness, then finding mates of intermediate genetic similarity maximizes reproductive success [4,18,19].

Since inbreeding and outbreeding affect fitness, it is logical to ask what processes mitigate these consequences in natural populations [20]. In the context of mating and genetic similarity, behavioral studies have focused on kin recognition and species recognition. While members of different species obviously tend to be poor mates [18,21], kin recognition can modulate the fitness effects of inbreeding and outbreeding [10] if mate preferences reflect fitness patterns. Species may have the innate ability to recognize kin [see discussions by 1,2,22], or that ability may have to be acquired with experience [1,21]. Learning processes are likely to play important roles in refining the ability to recognize kin, e.g. imprinting [21], phenotype matching [1] and individual familiarity [10,13,18,23]. Not surprisingly, researchers examining these processes typically use vertebrate species that live in defined social groups or have considerable parental care. However, hermaphroditic invertebrate systems that lack social organization and parental care afford an exciting opportunity to address questions of how mating patterns are influenced by both innate abilities and individual experience.

*Physa* is a genus of aquatic snails common throughout North America [24]. Physids are considered to be good dispersers and colonizers [25,26], and some species are globally invasive (e.g. *Physa acuta*) [27–29]. *Physa* live in diverse aquatic habitats ranging from large lakes and rivers to small streams, ditches, and temporary ponds [30]. Within these habitat types, population densities vary from hundreds of snails per m^2^ to fewer than one snail per m^2^ [31]. Given the high reproductive rates of physid snails, individuals may encounter relatives as potential mates, especially in smaller populations living in restricted habitats. In contrast, individuals may also encounter conspecifics from foreign populations as a result of transport via other organisms (e.g. water birds [25,32–35]) or abiotic factors (e.g. flooding [34]). Consequently, snails in some natural populations are likely to encounter potential mates with considerable genetic diversity while low diversity of mates may be encountered in other populations. Thus, recognition of genetic similarity may be an important selective pressure for physid snails in at least some natural populations. Interestingly, it is plausible that the general argument proposed for other systems (i.e. familiarity serves as a proximate mechanism for assessing kinship, where familiar individuals are likely to be relatives) may also hold true for snails. If snails have limited dispersal (e.g. restricted habitat) or have colonized new sites (small founding population), then individuals could have repeated interactions with each other. Koene and Ter Maat [36] and Häderer et al. [37] also discuss why familiarity, in contexts other than as a proxy for kin recognition, may or may not be important during mating interactions in different populations of aquatic snails.

Hermaphrodite snails are an excellent system for examining mating behaviors [20,38–42]. Since *Physa* are facultative simultaneous hermaphrodites, every mature individual, including itself, is a potential mate. *Physa* produce many offspring and mate readily, with easily observed mating behaviors [39,43]. For a successful copulation to occur, snails must physically contact each other, and then one snail assumes the male gender role by crawling onto the shell of the other snail, which becomes the prospective female by default. The snail acting as a male will position itself along the margin of the shell aperture of the prospective female by crawling in small circles on the female’s shell. Once in the proper alignment, the attempting male everts the penile complex (preputium); copulation occurs when the male’s preputium contacts the female’s gonopore and sperm is transferred. However, presumptive females can resist males through a variety of behaviors [39,43,44], including ‘biting’ behaviors where the female turns its head and the mouth region contacts the male’s preputium, presumably scraping it with the radula.

Pheromones likely play important role in attracting mates in many gastropods [45]. While the responses of physid snails to chemical cues associated with predation risk are well documented [46–57], they also respond to cues from uninjured conspecifics by increasing growth rates [58] and activity rates (unpublished data). Koene and Ter Maat [36] found that the presence of mucus trails influenced mate-choice decisions, but little other research has focused on possible roles of pheromones in mating interactions of aquatic pulmonate snails.

This study assessed whether previous experience with and the degree of relatedness between potential mates influenced behavior. I addressed these questions by observing the activity levels and mating interactions of aquatic snails in the laboratory. Previous work found that both close relatives (including self-fertilizing individuals) and very dissimilar partners produced fewer offspring relative to mates of intermediate similarity [44]. Thus, I predicted that mates of intermediate relatedness should be favored (i.e. greater escalation of interactions with less resistance) over both closely related and highly dissimilar mates. That is, snails would avoid extreme inbreeding and outbreeding. As in some snails [36] and vertebrate systems (e.g. naked mole-rats [59] and voles [60]), I also predicted that unfamiliar mates would generally be favored over familiar individuals. However, this predicted trend might be altered by conflicting preferences for degree of relatedness versus familiarity of potential mates. Finally, I predicted that previous experiences should also influence mating behavior with a current partner: previous exposure to preferred snails (in terms of relatedness or familiarity) should reduce motivation to mate with a current partner, whereas, previous exposure to non-preferred snails should increase mating with a current partner.

## Methods

### Rearing & setup

I collected *Physa gyrina* from two populations: a small stream in Jessamine County, KY, and from a small drainage ditch running parallel to train tracks near the University of Kentucky campus in Lexington, KY, USA. Collection sites were approximately 40 km apart. Wild-caught snails were used to initiate family lines in the laboratory. I isolated, measured and marked sexually mature, virgin snails (180 individuals) from the third laboratory-reared generation of family lines of the Jessamine County population. I assigned snails to five mate-type treatments (Fig 1: I.A): sibling pairs produced by a self-fertilizing parent (SS), sibling pairs from outcrossing parents (SO), cousin pairs (CN), unrelated intrapopulation pairs (UR), and interpopulation pairs (IP). Pairing third-generation laboratory-reared snails from the Jessamine County lines with first-generation laboratory-reared snails from the Lexington population formed IP pairs. Each relatedness treatment contained 20 replicate pairs.

**Fig 1.**
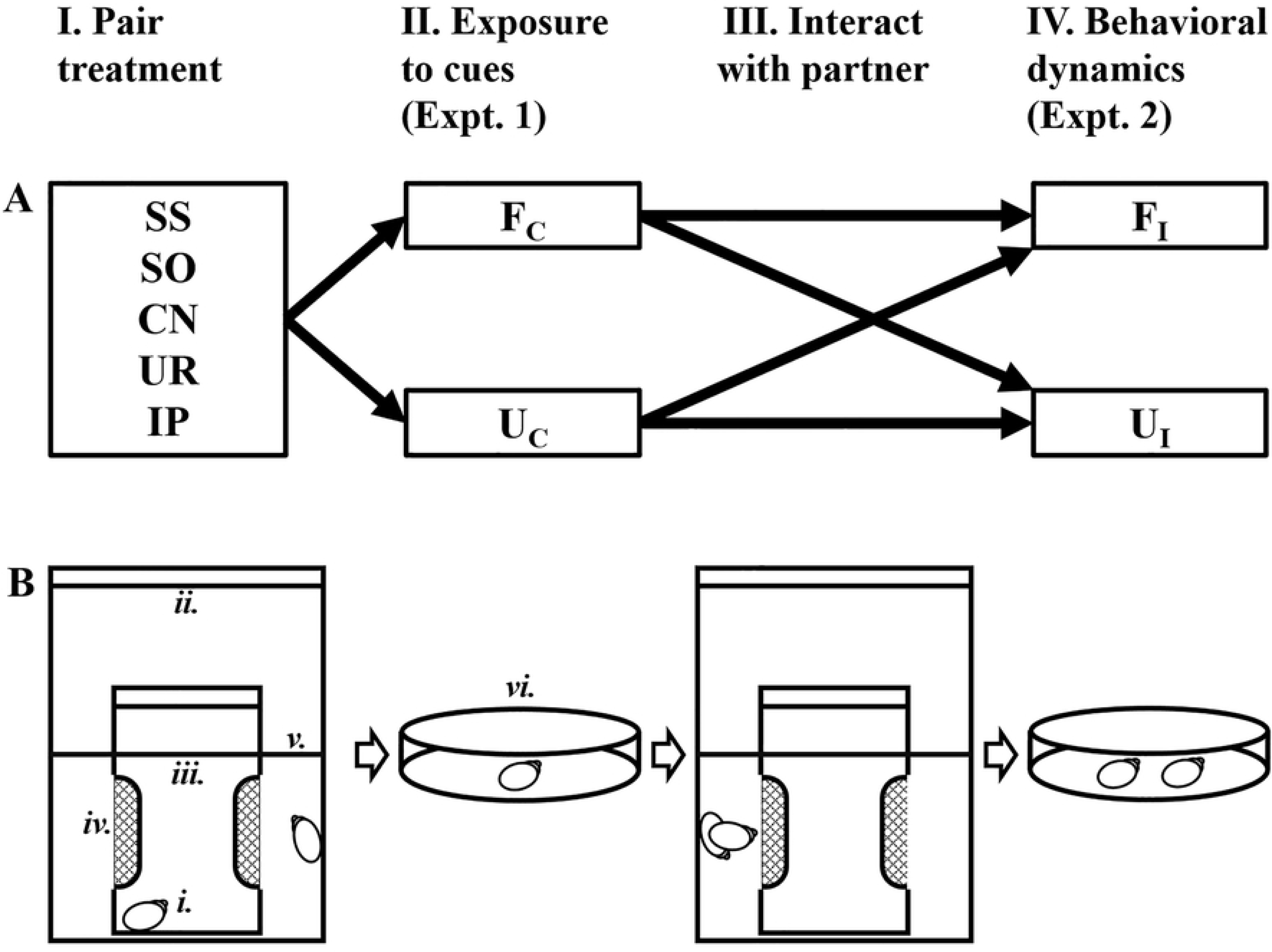
Diagram illustrating the experimental design. The top section (A) represents the 5×2×2 factorial design of experiment, while the bottom section (B) illustrates the pathways experienced by the snails during the study. Pairing treatments (I-A) were based on genetic similarity of partners (SS: sibling pairs from a selfed parent; SO: sibling pairs from outcrossed parents; CN: cousin pairs; UR: unrelated pairs; and IP: interpopulation pairs). Paired snails were exposed to each other’s chemical cues in a double-cup apparatus for two weeks (I-B: *i*. snail; *ii*. outer cup; *iii*. inner cup; *iv*. mesh window; *v*. waterline). Activity levels (II: Experiment 1) of individual snails were then recorded in Petri dishes (*vi*.) while exposed to either familiar (F_C_) or unfamiliar (U_C_) chemical cues. Next, partners were allowed to physically interact for 72 h (III). Last, behavioral dynamics (IV: Experiment 2) were observed during mating interactions between either familiar (F_I_) or unfamiliar (U_I_) individuals.

Paired snails were maintained together for two weeks so that individuals were exposed to chemical cues from their partner without being able to directly interact with each other. I placed each pair into an apparatus consisting of a smaller 250 ml plastic cup with two mesh windows resting inside a larger 900 ml plastic cup (Fig 1: I.B). One snail was placed in the small, inner cup while its partner was placed into the large, outer cup. Thus, partners were not able to physically interact but could indirectly assess each other by means of chemical cues passing through the mesh windows of the small cup. Lids on both cups prevented snails from interacting or escaping the apparatus. As described below, I used all snails in two successive experiments (Fig 1) in order to examine whether behavioral dynamics and mating interactions were influenced by the degree of relatedness between individuals, previous experience with the current partner, and exposure to cues from other potential mates. SYSTAT (Systat Software, San Jose, CA) was used for all statistical analyses with α = 0.05 as the threshold for significance.

### Experiment 1: Crawling rates & chemical cues

This stage of the experiment tested whether the locomotion rates of mature, virgin *P. gyrina* were influenced by either chemical familiarity or the degree of relatedness between individuals. Since mating propensity is associated with high locomotion rates in some gastropods [61], I predicted that crawling rates could serve as a proxy representing mate-search effort and would increase in response to chemical cues from unfamiliar snails and from snails of intermediate relatedness. To test these hypotheses, I exposed focal snails to chemical cues from conspecifics that varied in relatedness and prior chemical exposure (i.e. novel or familiar cues). I compared mean travel distances between treatments to determine whether snails differentially respond to conspecifics without direct interaction.

Snails from each of five relatedness categories were randomly assigned to a chemical-exposure treatment (Fig 1: II.). Snails in the chemically-familiar treatments were exposed to chemical cues from their own holding apparatus: snails from the inner cup received water drawn from the outer cup and vice versa. It should be noted that as a result of the mesh windows in the small cups, each snail was exposed to chemical cues from two potential sperm donors: themselves (because physids are simultaneous hermaphrodites) and the other snail in the apparatus. Snails assigned to the chemically-unfamiliar treatments were exposed to chemical cues generated by another pair of snails maintained in a different apparatus. Again, this exposed each focal snail to cues from two potential mates. In order to reduce confounding effects, each snail in the chemically-unfamiliar treatment was exposed to cues from an apparatus containing a pair that was related to the focal snail to the same degree as was its apparatus-partner. For example, a focal CN snail was exposed to cues from snails in an unfamiliar CN pair that were also cousins to the focal snail. Thus, the degree of relatedness remained constant while the familiarity changed.

I exposed each snail to the chemical cues by placing it in a Petri dish containing 50 ml of the appropriate treatment water. Snails’ paths were traced during a 3-minute observation period. Tracings were photographed, the images were digitized using ImageJ [62], and the travel distances were calculated for each snail. I compared crawl-distances between treatments, with shell length as a covariate (ANCOVAs). I also used orthogonal contrasts to specifically examine how genetic similarity influenced crawl-distances: a treatment group was compared to all more-related groups (i.e. IP versus all intrapopulation pairs; UR versus all related pairs; CN to all sibling pairs; SO to SS).

### Experiment 2: Behavior & mating interactions

Using the snails from experiment 1, I then tested whether behavioral dynamics during actual mating interactions were influenced by genetic relatedness, the exposure to chemical cues during experiment 1, and direct experience with the current partner. I predicted that all three factors would significantly affect the mating interactions: (i) there would be more resistance between sibling and interpopulation pairs [e.g. 44]; (ii) there would be greater escalations during interactions between sexually unfamiliar mates [e.g. 36], and (iii) previous exposure to chemical cues would alter interactions with the current partner. I predicted that previous exposure to cues from chemically unfamiliar snails would reduce motivation to mate with a current partner, while previous exposure only to cues from chemically familiar snails would increase the likelihood of mating with the current partner.

Following the first experiment, I returned the snails to their holding apparatus and allowed the partners to physically interact for 72 h (both snails were placed in outer cup; Fig 1: III.) since highly sexually motivated individuals (e.g. virgins) may not show strong mate preferences [8,63]. After the 72-h interaction period, I again isolated the individuals (one snail was returned to the inner cup) for 24 hours prior to testing. Pairs were then randomly re-assigned to a mating treatment (Fig 1: IV.) and, as in experiment 1, unfamiliar mates were equivalent to the snails’ familiar partners in terms of relatedness. Therefore, this was a 5 × 2 × 2 factorial design (relatedness of paired snails x exposure to familiar/unfamiliar chemical cue x interaction with familiar/unfamiliar partner). I placed paired snails in Petri dishes containing 50 ml of clean water and observed them for 60 minutes.

I quantified the behavioral dynamics observed during interactions by calculating mean conditional frequencies of escalation behaviors for each pair during the observation period as follows: number of contacts per hour, number of mountings per contact, number of positioning behaviors per mounting, number of preputium eversions per mounting, and number of copulations per preputium eversion. I also quantified gender-neutral, male-role, and female-role resistance and rejection responses observed during the interaction period by calculating mean conditional frequencies. The number of avoidance responses per number of contacts was considered a gender-neutral rejection response since neither snail attempted to occupy the male role. The frequency of rejections while occupying the male role was determined by the number of times males dismounted a female without attempting to copulate and without resistance behaviors from the female per mounting. ‘Biting’ behavior (mouth region contacts male’s preputium) was considered resistance to attempted copulations by the snail occupying the female role. Note that females need not resist unless a partner escalates its male function. If a snail prefers not to mate with a current partner, it should increase its resistance as the partner escalates. With preferred partners, a female’s resistance should not change with the partner’s escalation.

I compared behavioral dynamics between treatments with ANOVAs. ‘Mean sizes’ of and proportional ‘size differences’ between paired individuals were initially used as covariates but were dropped from the analyses since they did not significantly contribute to the models. Female-role resistance behaviors (‘bites’) were examined using linear regression analyses. Failure-time analyses [64] were used to compare the mean elapsed times until successful copulations occurred in each treatment. Failure-time analyses, which compare the rates at which interactions occur, may give an indication of mating preferences that are not reflected in the conditional frequencies of behaviors described above. That is, a treatment factor may influence the speed at which behaviors occur and interactions escalate without affecting the frequencies of those behaviors.

## Results

### Experiment 1: Crawling rates & chemical cues

The snails’ activity levels (crawl-distances) were significantly influenced by the relatedness of their partners (Fig 2; ANCOVA: F_4,142_ = 4.77, p = 0.001). Orthogonal contrasts showed that within intrapopulation pairs, activity decreased with increasing relatedness of partners. Snails crawled farther when exposed to cues from: 1) UR compared to related snails, 2) CN compared to siblings, and 3) SO compared to SS. Crawling distances were not affected by familiarity treatments (F_1,142_ = 0.29, p = 0.59; mean distance ± SE (cm): familiar treatment = 21.19 ± 0.75, unfamiliar treatment = 21.31 ± 0.83) or body size (F_1,142_ = 2.72, p = 0.10). There was no significant interaction between pairing and familiarity treatments (F_4,142_ = 2.08, p = 0.09).

**Fig 2.**
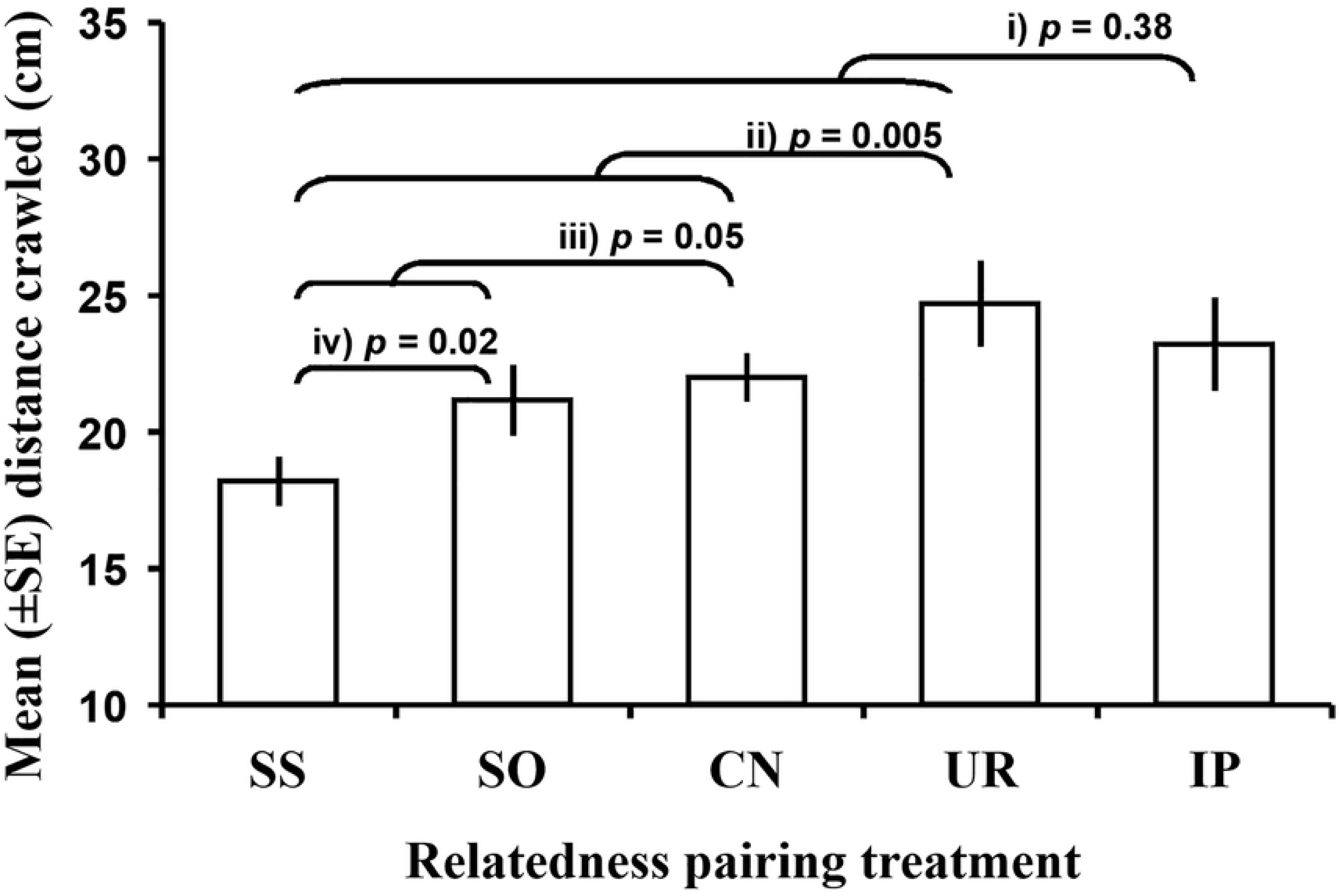
Mean (±SE) crawling distances (cm) of snails in each treatment. (SS: sibling pairs from a selfed parent; SO: sibling pairs from outcrossed parents; CN: cousin pairs; UR: unrelated pairs; and IP: interpopulation pairs) Orthogonal contrasts compared *i*) IP versus all intrapopulation pairs, *ii*) UR versus all related pairs, *iii*) CN versus all sibling pairs, and *iv*) SO versus SS.

### Experiment 2: Behavior & mating interactions

The degree of relatedness between snails did not influence the interaction outcome (Table 1), the latency of copulation (failure-time analysis: χ^2^ = 3.90, df = 4, *p* = 0.42), the escalation of male-role behaviors (Table 2) or male-role rejection frequencies (Table 3) during interactions. In contrast, female-role resistance varied with the relatedness of the partner. Female-role biting behaviors (female’s mouth region contacts male’s preputium) significantly increased with male mating attempts (preputium eversion frequencies) for both highly related SS snails and dissimilar IP snails (Table 4A). Conversely, snails did not significantly increase female-role resistance biting behaviors against highly motivated males with more intermediate degrees of relatedness (SO, CN, and UR snails; Table 4A).

**Table 1.**
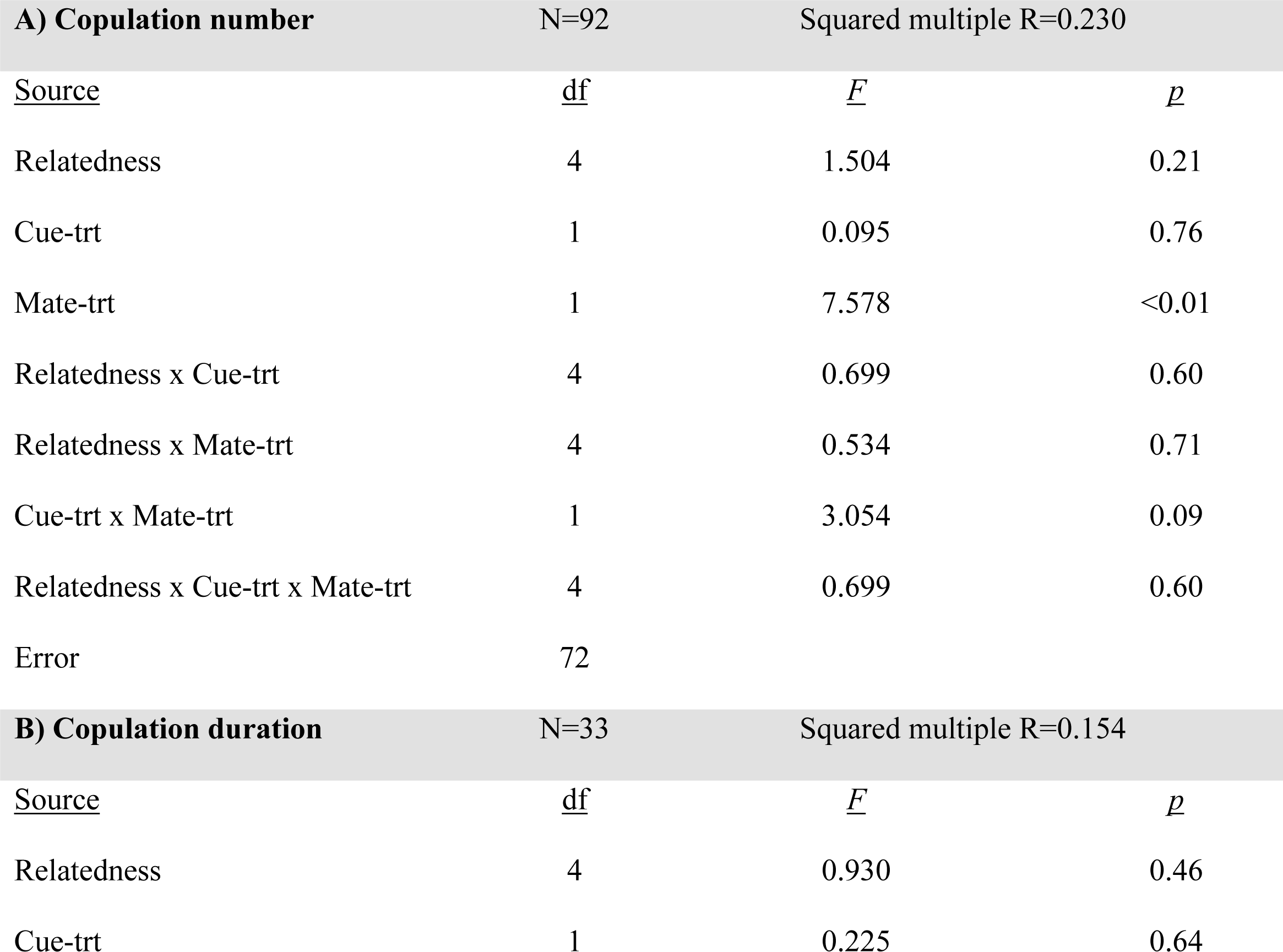
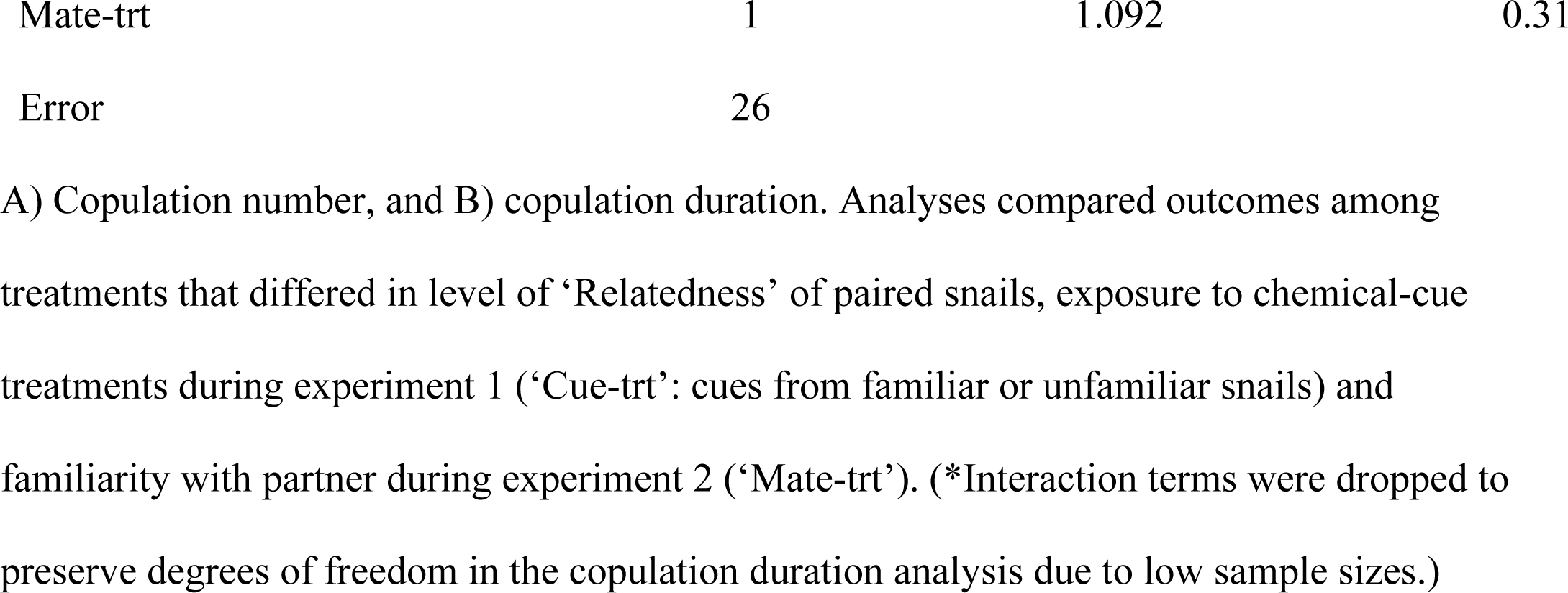
Results of ANOVAs testing for treatment effects on the outcomes of mating interactions between paired snails (*Physa gyrina*).

| **A) Copulation number** |  | N=92 |  | Squared multiple R=0.230 |  |
| --- | --- | --- | --- | --- | --- |
| <u>Source</u> | <u>df</u> |  | <u>F</u> |  | <u>p</u> |
| Relatedness | 4 |  | 1.504 |  | 0.21 |
| Cue-trt | 1 |  | 0.095 |  | 0.76 |
| Mate-trt | 1 |  | 7.578 |  | <0.01 |
| Relatedness x Cue-trt | 4 |  | 0.699 |  | 0.60 |
| Relatedness x Mate-trt | 4 |  | 0.534 |  | 0.71 |
| Cue-trt x Mate-trt | 1 |  | 3.054 |  | 0.09 |
| Relatedness x Cue-trt x Mate-trt | 4 |  | 0.699 |  | 0.60 |
| Error | 72 |  |  |  |  |
| **B) Copulation duration** |  | N=33 |  | Squared multiple R=0.154 |  |
| <u>Source</u> | <u>df</u> |  | <u>F</u> |  | <u>p</u> |
| Relatedness | 4 |  | 0.930 |  | 0.46 |
| Cue-trt | 1 |  | 0.225 |  | 0.64 |

|  |  |  |  |
| --- | --- | --- | --- |
| Mate-trt | 1 | 1.092 | 0.31 |
| Error | 26 |  |  |

A) Copulation number, and B) copulation duration. Analyses compared outcomes among treatments that differed in level of ‘Relatedness’ of paired snails, exposure to chemical-cue treatments during experiment 1 (‘Cue-trt’: cues from familiar or unfamiliar snails) and familiarity with partner during experiment 2 (‘Mate-trt’). (\*Interaction terms were dropped to preserve degrees of freedom in the copulation duration analysis due to low sample sizes.)

**Table 2.**
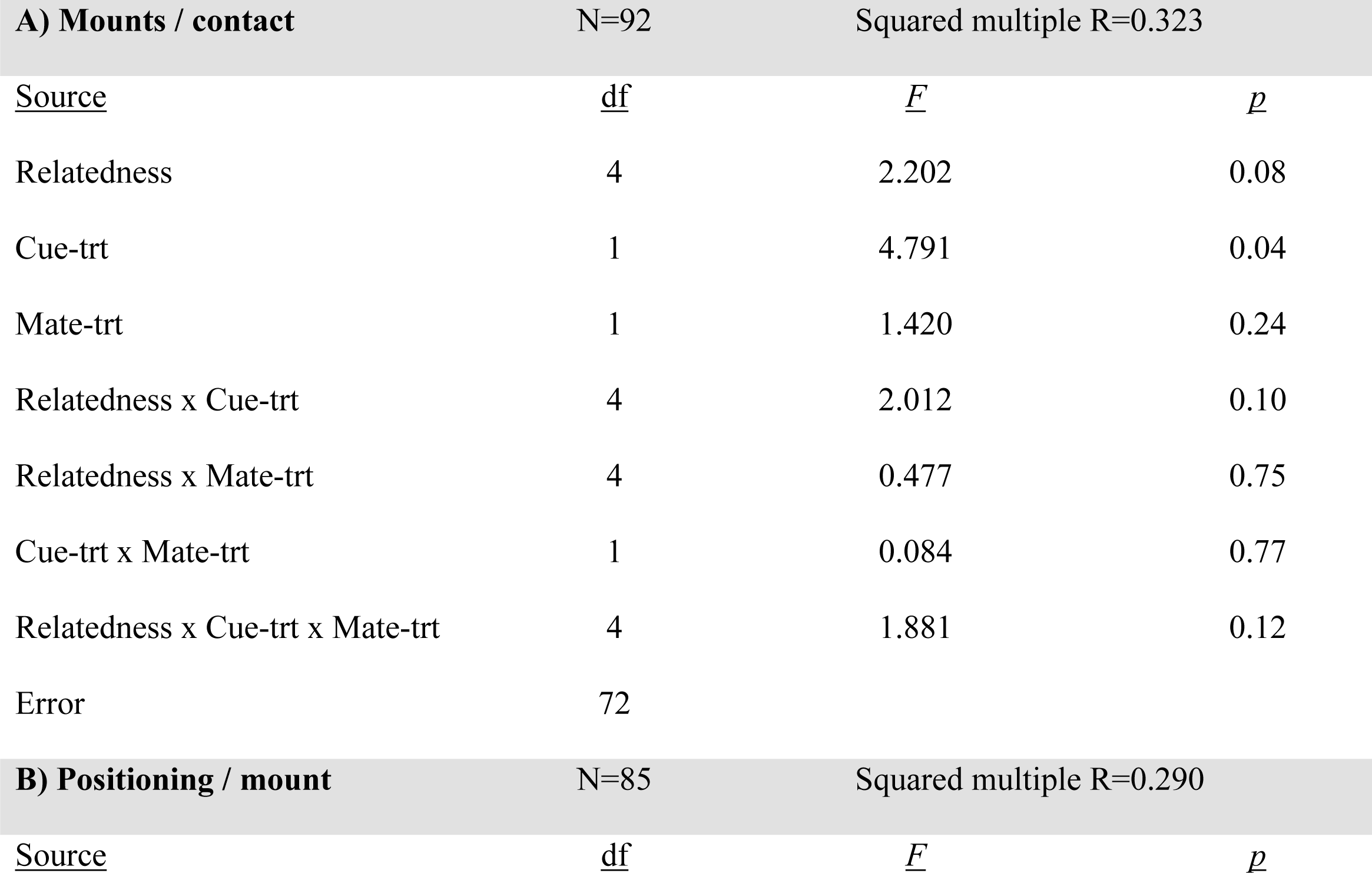
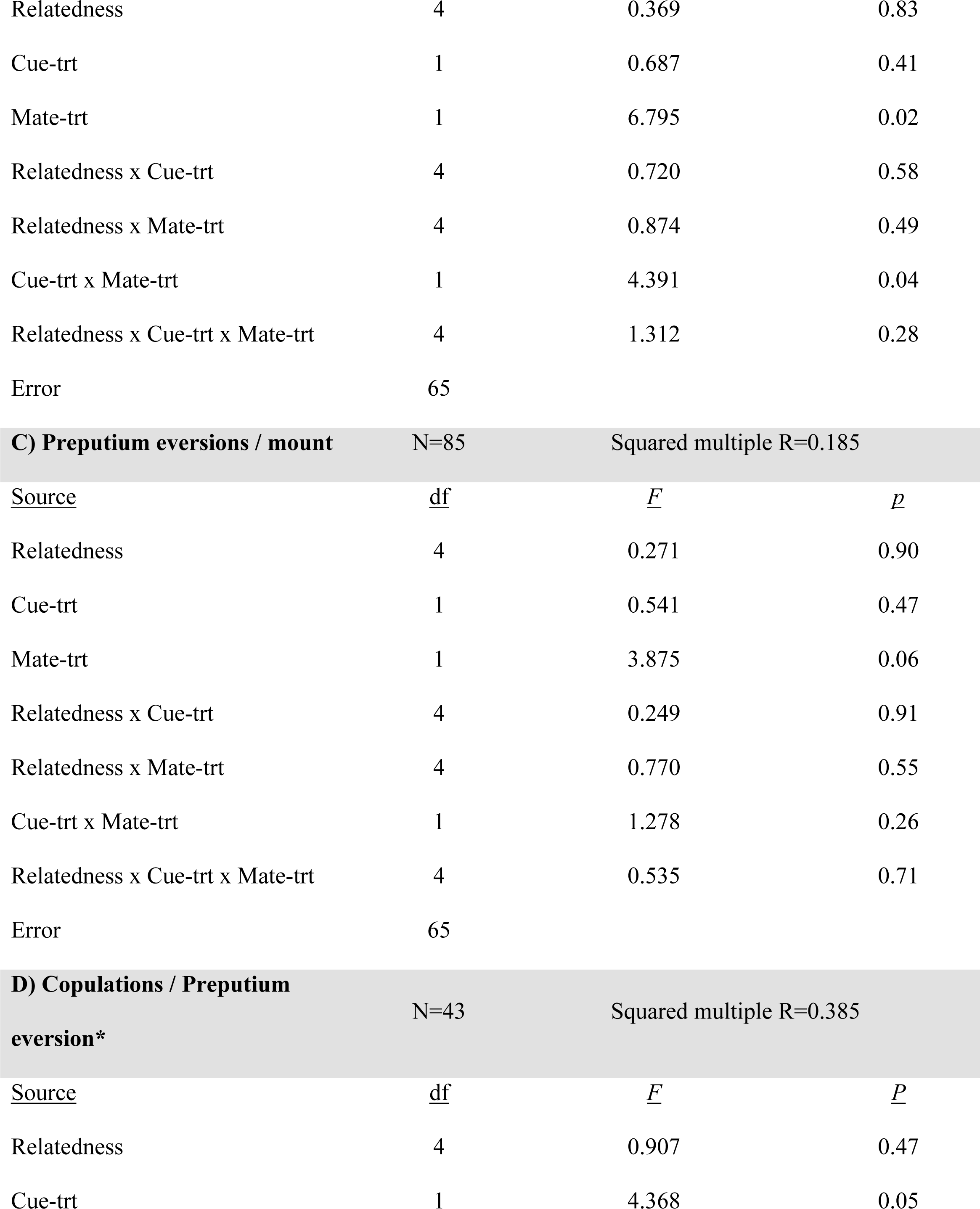
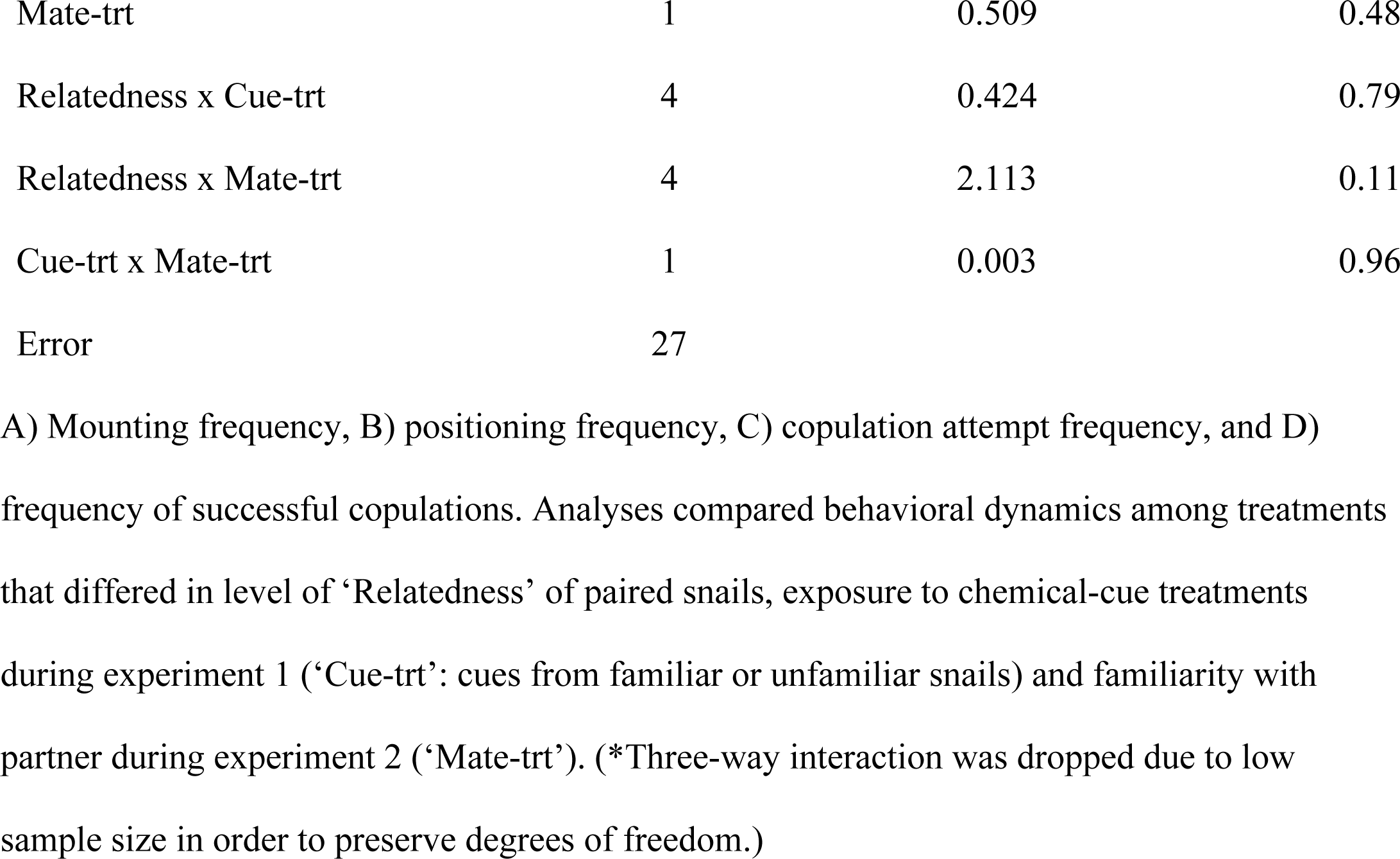
Results of ANOVAs testing for treatment effects on various behavioral aspects of mating interactions between paired snails (*Physa gyrina*).

**Table 3.**
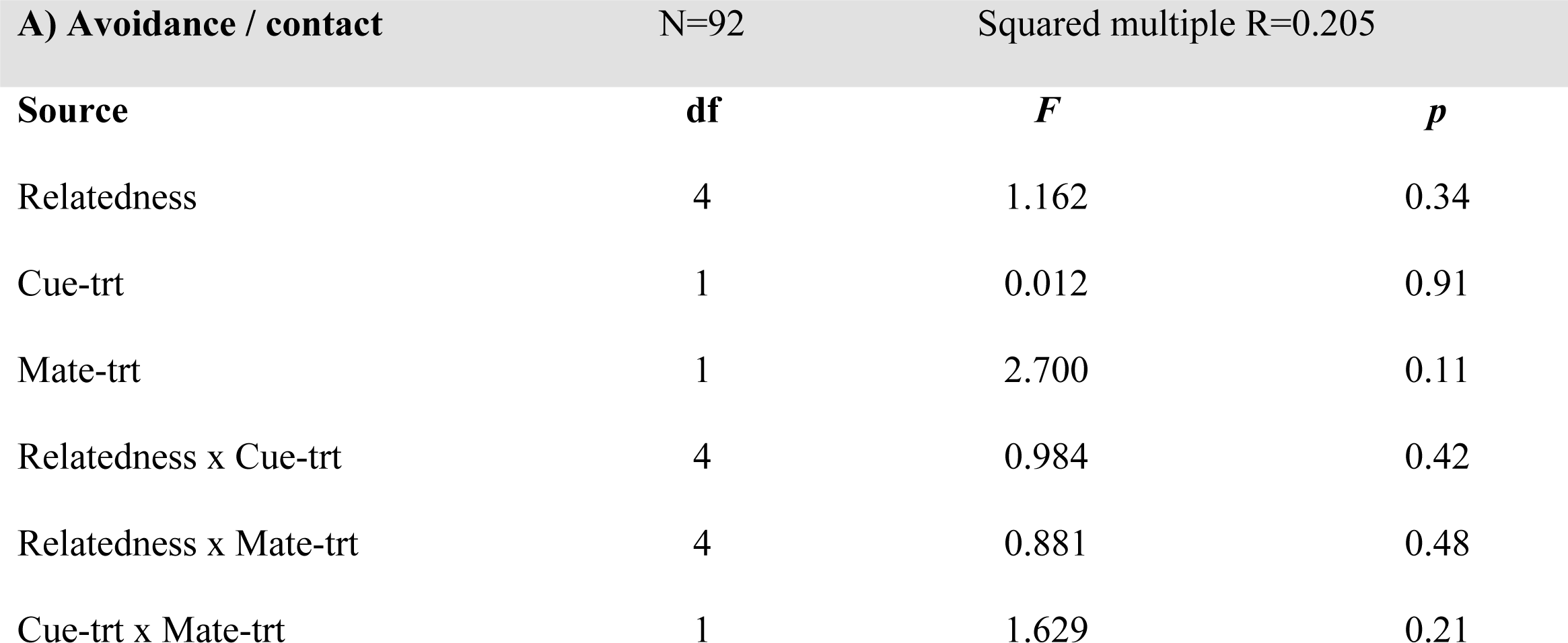
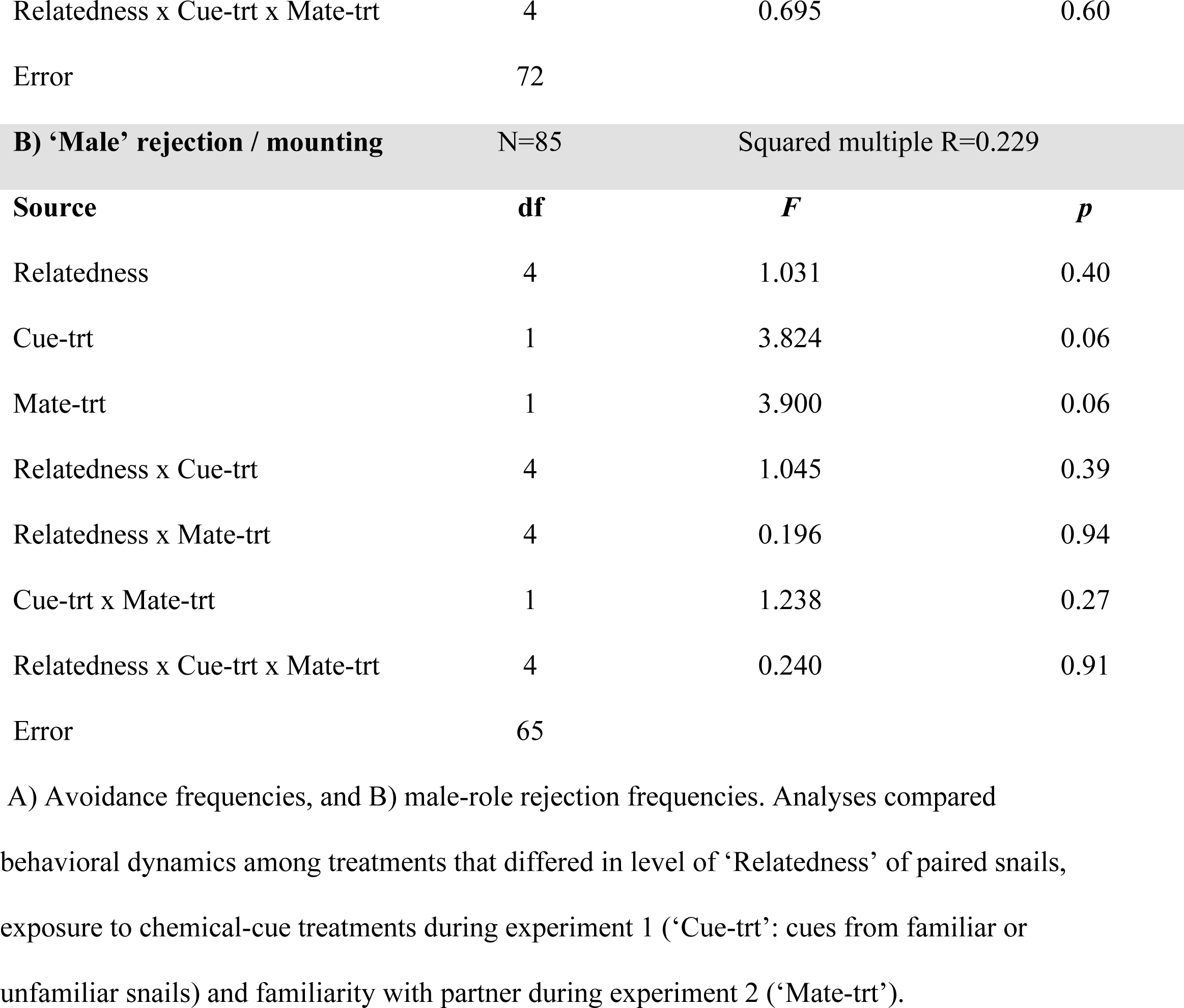
Results of ANOVAs testing for treatment effects on mate-rejection behaviors during mating interactions between paired snails (*Physa gyrina*)

| A) Avoidance / contact |  | N=92 |  | Squared multiple R=0.205 |
| --- | --- | --- | --- | --- |
| Source | df | F |  | p |
| Relatedness | 4 | 1.162 |  | 0.34 |
| Cue-trt | 1 | 0.012 |  | 0.91 |
| Mate-trt | 1 | 2.700 |  | 0.11 |
| Relatedness x Cue-trt | 4 | 0.984 |  | 0.42 |
| Relatedness x Mate-trt | 4 | 0.881 |  | 0.48 |
| Cue-trt x Mate-trt | 1 | 1.629 |  | 0.21 |

|  |  |  |  |
| --- | --- | --- | --- |
| Relatedness x Cue-trt x Mate-trt | 4 | 0.695 | 0.60 |
| Error | 72 |  |  |

| B) 'Male' rejection / mounting |  | N=85 | Squared multiple R=0.229 |
| --- | --- | --- | --- |
| Source | df | F | p |
| Relatedness | 4 | 1.031 | 0.40 |
| Cue-trt | 1 | 3.824 | 0.06 |
| Mate-trt | 1 | 3.900 | 0.06 |
| Relatedness x Cue-trt | 4 | 1.045 | 0.39 |
| Relatedness x Mate-trt | 4 | 0.196 | 0.94 |
| Cue-trt x Mate-trt | 1 | 1.238 | 0.27 |
| Relatedness x Cue-trt x Mate-trt | 4 | 0.240 | 0.91 |
| Error | 65 |  |  |

A) Avoidance frequencies, and B) male-role rejection frequencies. Analyses compared behavioral dynamics among treatments that differed in level of 'Relatedness' of paired snails, exposure to chemical-cue treatments during experiment 1 ('Cue-trt': cues from familiar or unfamiliar snails) and familiarity with partner during experiment 2 ('Mate-trt').

**Table 4.**
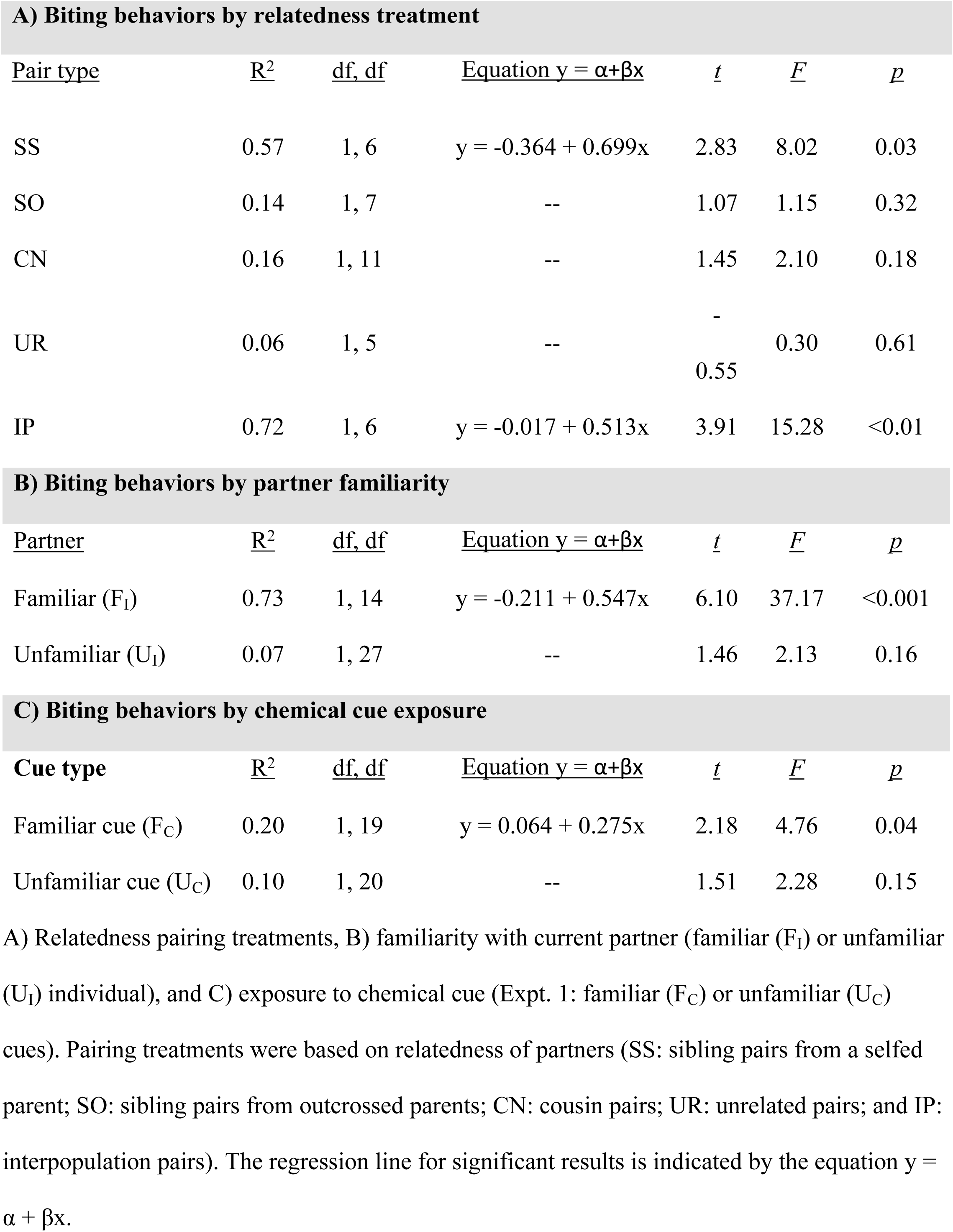
Regression analyses comparing female-role resistance behaviors following partner’s male-role preputium eversions during mating interactions between paired snails (*Physa gyrina*).

| A) Biting behaviors by relatedness treatment |  |  |  |  |  |  |
| --- | --- | --- | --- | --- | --- | --- |
| Pair type | R<sup>2</sup> | df, df | Equation $y = \alpha + \beta x$ | $t$ | $F$ | $p$ |
| SS | 0.57 | 1, 6 | $y = -0.364 + 0.699x$ | 2.83 | 8.02 | 0.03 |
| SO | 0.14 | 1, 7 | -- | 1.07 | 1.15 | 0.32 |
| CN | 0.16 | 1, 11 | -- | 1.45 | 2.10 | 0.18 |
| UR | 0.06 | 1, 5 | -- | -<br>0.55 | 0.30 | 0.61 |
| IP | 0.72 | 1, 6 | $y = -0.017 + 0.513x$ | 3.91 | 15.28 | <0.01 |

B) Biting behaviors by partner familiarity
| <u>Partner</u> | <u>R<sup>2</sup></u> | <u>df, df</u> | <u>Equation $y = \alpha + \beta x$</u> | <u>t</u> | <u>F</u> | <u>p</u> |
| --- | --- | --- | --- | --- | --- | --- |
| Familiar (F<sub>I</sub>) | 0.73 | 1, 14 | $y = -0.211 + 0.547x$ | 6.10 | 37.17 | <0.001 |
| Unfamiliar (U<sub>I</sub>) | 0.07 | 1, 27 | -- | 1.46 | 2.13 | 0.16 |

C) Biting behaviors by chemical cue exposure
| <u>Cue type</u> | <u>R<sup>2</sup></u> | <u>df, df</u> | <u>Equation $y = \alpha + \beta x$</u> | <u>t</u> | <u>F</u> | <u>p</u> |
| --- | --- | --- | --- | --- | --- | --- |
| Familiar cue (F<sub>C</sub>) | 0.20 | 1, 19 | $y = 0.064 + 0.275x$ | 2.18 | 4.76 | 0.04 |
| Unfamiliar cue (U<sub>C</sub>) | 0.10 | 1, 20 | -- | 1.51 | 2.28 | 0.15 |

A) Relatedness pairing treatments, B) familiarity with current partner (familiar (F<sub>I</sub>) or unfamiliar (U<sub>I</sub>) individual), and C) exposure to chemical cue (Expt. 1: familiar (F<sub>C</sub>) or unfamiliar (U<sub>C</sub>) cues). Pairing treatments were based on relatedness of partners (SS: sibling pairs from a selfed parent; SO: sibling pairs from outcrossed parents; CN: cousin pairs; UR: unrelated pairs; and IP: interpopulation pairs). The regression line for significant results is indicated by the equation $y = \alpha + \beta x$ .

Mating interactions were significantly influenced by prior interactions with the current partner. Snails that had previous interactions with each other had fewer copulations than unfamiliar mates (‘Mate-trt’ in Table 1; mean number of copulations per pair ± SE: familiar snails = 0.31 ± 0.08, unfamiliar snails = 0.68 ± 0.12). Unfamiliar snails also mated sooner than familiar partners (Fig 3; failure-time analysis: χ^2^ = 4.60, df = 1, *p* = 0.03). These differences reflect changes in both male-role and female-role behaviors; there were lower frequencies of male-role escalation behaviors and increasing female-role resistance during mating interactions between familiar snails. While the analyses comparing male-role rejection and preputium eversion frequencies between familiarity treatments were marginally nonsignificant (Tables 2 & 3), the trends suggest that snails occupying the male role during an interaction with an unfamiliar partner were less likely to reject the partner (mean male-role rejection frequencies ± SE: familiar snails = 0.70 ± 0.05, unfamiliar snails = 0.57 ± 0.05) and were more likely to attempt a copulation (mean preputium eversion frequencies ± SE: familiar snails = 0.33 ± 0.10, unfamiliar snails = 0.78 ± 0.17). Snails occupying the female-role increased their resistance to highly motivated, familiar partners but did not significantly increase their resistance to highly motivated, unfamiliar partners (Table 4B).

**Fig 3.**
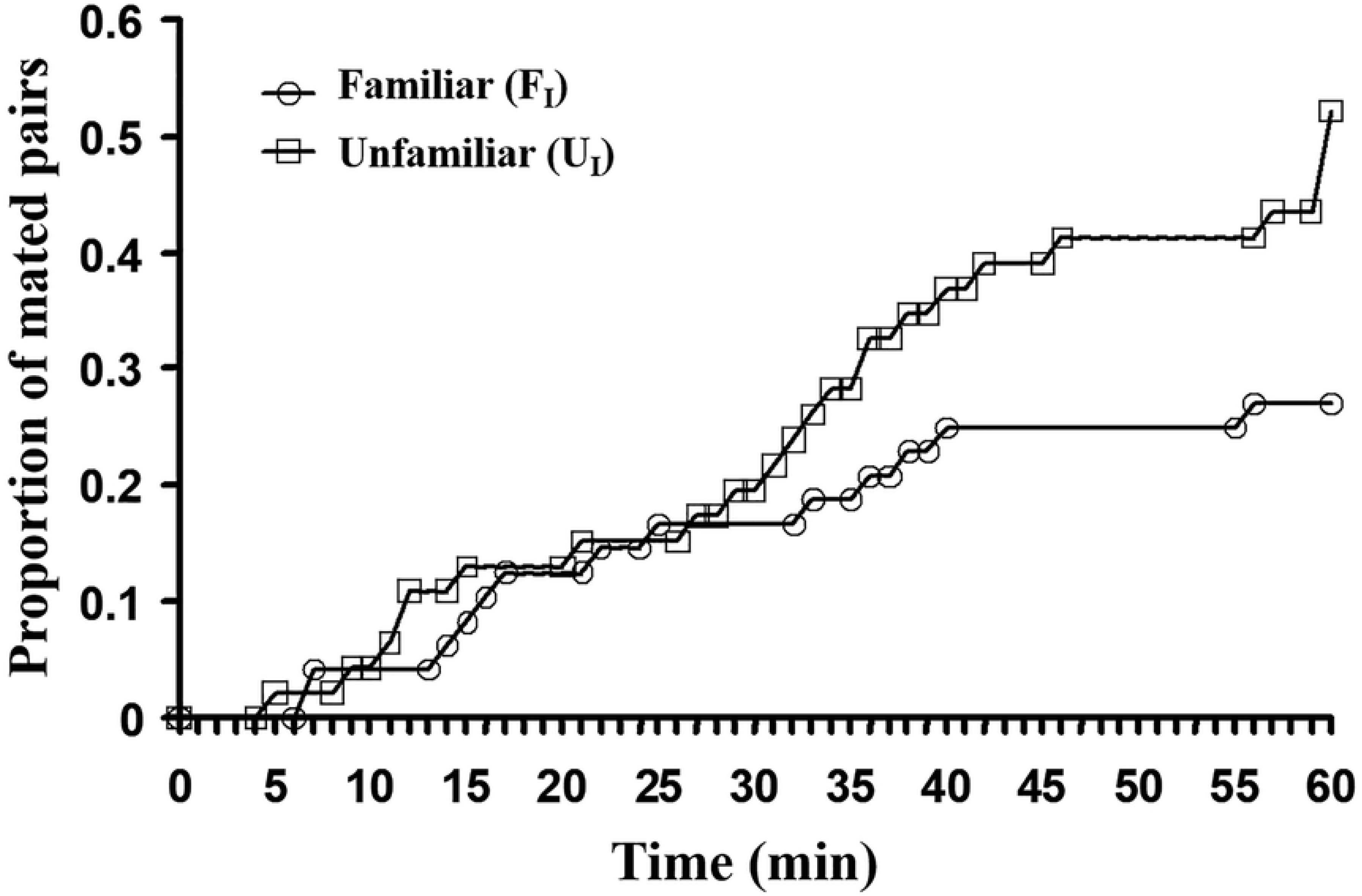
Mating distribution functions. Proportions of pairs of snails that mated when interacting with a familiar (F_I_) or an unfamiliar (U_I_) partner over time.

The chemical-cue familiarity treatments that snails experienced in the first experiment also significantly affected the mating interactions observed in the second experiment. Individuals that were exposed to chemical cues from unfamiliar snails in the first experiment were more likely to mount their partners (‘Cue-trt’ in Table 2A; mean mounting frequencies ± SE: exposed to familiar cues = 0.47 ± 0.04, exposed to unfamiliar cues = 0.62 ± 0.05), but also tended to reject the female more often during an interaction (‘Cue-trt’ in Table 3B – marginally nonsignificant; mean ‘male’ rejection frequencies ± SE: exposed to familiar cues = 0.58 ± 0.06, exposed to unfamiliar cues = 0.70 ± 0.04). Positioning frequencies were affected by a significant interaction between the familiarity treatments of the two experiments (Table 2B; Fig 4). Bonferroni *post hoc* tests indicated that snails exposed to familiar cues during the first experiment significantly increased positioning frequencies when interacting with an unfamiliar individual. Although there was no difference in the latency to mating (failure-time analysis: χ^2^ = 0.16, df = 1, p = 0.69), copulation frequencies were greater for snails that had experienced familiar chemical cues during the first experiment (Table 2D; mean copulation frequencies ± SE: exposed to familiar cues = 0.65 ± 0.09, exposed to unfamiliar cues = 0.50 ± 0.08). Snails occupying the female role that had been exposed to familiar chemical cues during the first experiment resisted highly motivated males during interactions, but this was not the case for snails that had been exposed to unfamiliar cues in the first experiment (Table 4C).

**Fig 4.**
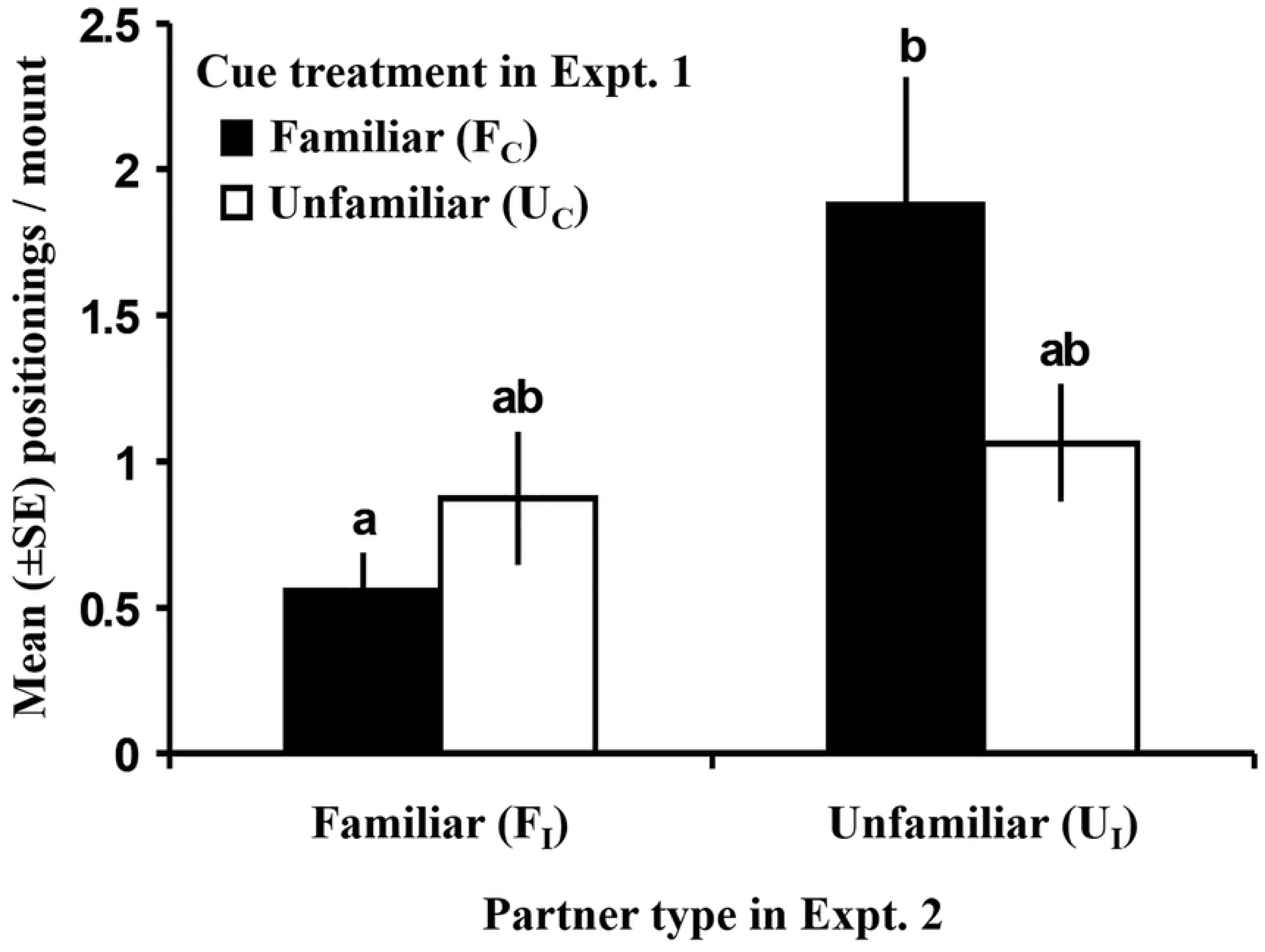
Mean number (±SE) of positioning behaviors per mounting between paired individuals during mating interactions. In Experiment 1, individuals were exposed to cues from either chemically familiar (F_C_) or unfamiliar (U_C_) snails for a 3-min period. Snails then interacted with either a physically familiar (F_I_) or unfamiliar (U_I_) partner in Experiment 2. Bars sharing the same letter are not statistically different.

Neither a snail’s size nor its crawl-distance (first experiment) was related to the individual’s conditional frequencies of behaviors during mating interactions (Table 5). Moreover, there was no trend for size-based gender occupation during successful copulations (mean size (mm) ± SE: females = 7.25 ± 0.04, males = 7.23 ± 0.05; paired t-test: *t*_33_ = 0.14, *p* = 0.89).

**Table 5.** Results of multiple regression analyses testing for relationships of body size (Size) and activity levels (Distance) of *Physa gyrina* with various behavioral aspects of mating interactions.

| Conditional frequencies | <u>Adjusted squared</u><br><u>multiple R</u> | <u>Size</u> |  | <u>Distance</u> |  |
| --- | --- | --- | --- | --- | --- |
|  |  | <u>t</u> | <u>p</u> | <u>t</u> | <u>p</u> |
| Mounts / contact | 0.01 | 1.39 | 0.17 | 0.17 | 0.87 |
| Positioning / mount | 0.03 | 1.38 | 0.17 | 1.61 | 0.11 |
| 'Male rejection' / mounting | 0.03 | 1.51 | 0.14 | 1.54 | 0.13 |
| Preputium eversions / mount | 0.01 | 1.24 | 0.22 | 1.14 | 0.26 |
| Copulations / preputium eversion | 0.01 | 0.10 | 0.92 | 0.09 | 0.93 |

## Discussion

This study indicates that the degree of relatedness between individuals and the contexts in which mating interactions occur have important implications for mating systems. As a whole, the results suggest that potential mates are being assessed and treated differently. Two lines of evidence supported the prediction that there would be behavioral differences between pairs with different degrees of relatedness. First, snails crawled significantly farther in response to chemical cues from UR snails than to related snails. Second, snails occupying the female role during mating interactions showed greater resistance toward their partners in both highly inbred and outbred pairings. Behaviors of snails occupying the male role tended not to differ between relatedness treatments. The results also supported the prediction that unfamiliar partners would be preferred over familiar partners: unfamiliar pairings had more matings, shorter latency times to matings, less female-role resistance behavior, and, perhaps, less escalation of male-role behaviors. Previous exposure to chemical cues also significantly affected subsequent mating encounters. Although there was no evidence for any interactions between relatedness and individual familiarity, the data suggest that the interplay between familiarity with a current potential partner and an individual’s previous experiences may be important factors in mate choice.

Kin recognition (or discrimination) may be defined as the differential treatment of conspecifics based on their degree of genetic relatedness [22]. In a reproductive context, kin recognition requires individuals to assess the degree of relatedness of potential mates [14] and base their mating strategies on those assessments. Accordingly, selection pressures should act to minimize ‘acceptance errors’ (accepting low-quality mates) and ‘rejection errors’ (rejecting high-quality mates; terminology of Sherman et al. [22]). Thus, if fitness consequences vary with the relatedness of mates, then kin discrimination should reduce the frequency of ‘errors’ and help maximize reproductive success.

A previous study demonstrated that sibling and interpopulation pairs suffered reduced reproductive success [44]. Thus, the prediction for this study was that the snails would prefer cousins with intermediate relatedness [10,11,19,65], while avoiding sibling (inbreeding avoidance [17,66–69]) and interpopulation matings (outbreeding avoidance [20,70]). However, complex interactions arise when one individual possesses both genders [38,71], and preference strength may depend on numerous factors, such as, energy allocation, opportunity costs, and recent interactions. The costs of misallocating energy and effort toward low quality mates should usually make females choosier than males [18,72,73]. Before engaging in a mating interaction (and thereby assuming a gender role; experiment 1), snails increased activity levels with decreasing degrees of relatedness with their potential mates (differing from previous results [44]). Fearnley [61]argued that mating propensity is associated with high locomotion rates in some gastropods. This suggests that the snails were either avoiding inbred matings by decreasing activity in response to close relatives, or they increased search efforts for less related mates. While snails acting as males exhibited few preferences for mates based on relatedness, those acting in the female role significantly increased resistance to highly motivated SS and IP males. Thus, one interpretation for the lower crawling rates in response to cues from close relatives, given the gender-role specific strategies, is that reduced activity is a mechanism to avoid fitness costs should that individual become ‘female’ during an interaction. Consequently, the snails’ gender-specific behavioral strategies (no discrimination when acting as a male, but alter resistance levels to mates when acting as a female) should maximize individual fitness.

The potential roles of previous experience and individual familiarity during mating interactions have received considerable attention, typically in vertebrate systems that are philopatric, have extensive parental care, or live in social groups. The crux of the argument is often that familiarity serves as a proximate mechanism for assessing kinship, where familiar individuals are likely to be relatives. Thus, a focal animal can avoid inbreeding by preferring unfamiliar individuals as mates [e.g. 59,60]. The results of this study are consistent with this hypothesis as the snails preferred unfamiliar mates during interactions. However, it is difficult to envision the utility of recognizing individuals in a high-density population of a non-social species that lacks parental care: why would this be adaptive? Plausible mechanisms aside from individual recognition per se can also explain the observed preference patterns exhibited by the snails. One might expect familiarity to arise in species that regularly occur in relatively small populations living in patchily distributed or spatially restricted areas (e.g. small streams) where individuals may repeatedly interact with each other. Individuals might become habituated to the cues produced by regularly encountered conspecifics. This would likely increase the frequency of ‘rejection errors’, where high-quality mates are rejected because they are familiar. Individuals that are unfamiliar produce cues that may be perceived as novel and stimulating, thereby eliciting more intense responses than familiar individuals (e.g. Coolidge effect [36]). This could consequently increase ‘acceptance errors,’ where low-quality, unfamiliar mates are accepted over familiar, high-quality mates. In this study it is possible that variations in water quality between holding apparatuses might have influenced some aspect of the phenotype, thereby creating perceived differences between treatment groups. However, since every apparatus received the same source water and food, it seems unlikely that their water qualities would diverge to the extent that the observed behavioral patterns would consistently arise.

The effects of a 3-minute exposure to chemical cues on behaviors during mating interactions 96 hours later were intriguing. The results suggest that an individual’s previous experiences can strongly influence its subsequent mate-choice decisions. Cross-fostering studies with juvenile animals have demonstrated the potential long-term effects of experience on mating behaviors in species with parental care [60,74,75]. The present study shows that even brief and indirect interactions between sexually mature individuals have demonstrable and lasting effects on mating behaviors. Additionally, comparing the crawling-rate patterns in this experiment to previous results suggests that experience with another snail (indirect assessment within apparatus) influenced behaviors related to search effort. Naïve snails (having had no opportunity to assess conspecifics) exhibited no pattern related to the genetic similarity of potential mates [44], but snails that had an opportunity to assess a conspecific (current study) exhibited significant differences in crawling rates between relatedness treatments.

This type of system may be useful in addressing how mate preferences are formed and how potential mates are assessed, e.g. comparing “best-of-*n*,” “rare-male”, threshold, and sequential sampling models [76–79]. Future experiments could test predictions about how the encounter frequencies of potential types of mates in a population would influence mating behaviors. For instance, if unfamiliar individuals are preferred, then previous exposure to unfamiliar partners (or high frequencies of unfamiliar individuals) should make individuals choosier.

It is worth noting that body size had no effect on mating interactions during this experiment, but strongly influenced behaviors in other experiments [39,43,44] - also see [80]. Two possible explanations for this discrepancy are that the actual size differences between partners were relatively minor, and that the influence of familiarity and experience took precedence over size differences. That is, the preferences for unfamiliar individuals may overwhelm the effects of body size on behavioral dynamics.

This study demonstrates that relatedness, familiarity with a current potential partner, and prior experiences influence behavioral dynamics during mating interactions. These findings imply that snails can discriminate between potential mates, and suggests a number of areas for future research that are applicable to both hermaphroditic and dioecious systems. For a better understanding of mating patterns seen in nature, we need more information about what kinds of individuals are encountered as potential mates and about the genetic variance within and between populations. Future studies should continue to consider multiple mechanisms underlying mate preferences, as well as, possible interactions between these mechanisms. In addition to considering behavioral asymmetries between males and females in dioecious species, theoretical and empirical work should continue to examine the prevalence of behavioral gender-specific mating preferences in simultaneous hermaphrodites. While these tasks pose a significant challenge, the results will increase our comprehension of mating patterns observed in natural systems.

## Acknowledgments

Discussion and comments by A. Sih, P. Crowley, D. Westneat, R. C. Sargent, C. Akins, C. Fox, and E. McCarthy greatly improved this work. The help and insights provided by all of the members of the Sih, Crowley and Westneat lab groups were greatly appreciated. E. McCarthy and W. Weng aided in raising and maintaining the snail lines. This paper was based on dissertation work submitted in partial fulfilment of the requirements for a Ph.D. from the University of Kentucky.

## Supporting information

**S1 Table. Raw data of activity levels from Experiment 1**.

**S2 Table. Raw data of behavioral dynamics during interactions from Experiment 2**.

